# The inner dynamics of positive human-animal interactions: investigating the roles of oxytocin, opioids, dopamine, serotonin and the proteome

**DOI:** 10.1101/2025.07.24.666545

**Authors:** Oceane Schmitt, Suzanne Truong, Winfried Otten, Hana Volkmann, Julia Greiner, Marlies Dolezal, Karin Hummel, Attilio Rocchi, Jean-Loup Rault

## Abstract

Research on the neuroendocrine basis of positive interactions has predominantly focused on oxytocin (OT), although dopamine (DA) and opioids also play crucial roles. Furthermore, these neurotransmitters are known to interact with each other but have seldom been studied concurrently. In this study, we quantified longitudinal changes in these neurotransmitters using a within-subject, 2 × 2 factorial design by varying human familiarity (familiar versus unfamiliar) and contact type (positive contacts versus ignoring) for 10 min human-pig interaction sessions. We repeatedly sampled cerebrospinal fluid from 10 pigs through a spinal catheter 65 and 5 min before the test and at 10, 30, 60, 120 and 240 min after the start of the test. Samples at various timepoints were analysed for OT, DA metabolites 3,4-dihydroxyphenylacetic acid (DOPAC) and homovanillic acid (HVA), serotonin metabolite 5-hydroxyindoleacetic acid (5-HIAA), β-endorphin opioids concentrations, and using proteomics to explore novel protein candidates. The test condition (human familiarity × contact type) had a significant effect on the concentration of β-endorphin (F_1,118.59_ = 4.45; P = 0.04) and 5-HIAA (F_1,73.47_ = 5.02; P = 0.03), and tended to affect the concentration of oxytocin (F_1, 119.8_ = 3.19; P = 0.08) and DOPAC (F_1,70.57_ = 3.31; P = 0.07), but pair-wise comparisons were not significant. There was only a minor effect on the pigs’ behaviour, which suggests that the test conditions may have had limited effect. Nevertheless, this approach provides a valuable method to study neurotransmitter changes over time and simultaneously.

**Highlights:**

- Oxytocin was higher when pigs were ignored than when they had positive contacts
- Oxytocin was highest in the novel condition of an unfamiliar human ignoring the pig
- Human familiarity and contact type affected β-endorphin and 5-HIAA concentrations
- Dopamine metabolites and the proteome were not affected by the test conditions

## 1. Introduction

Positive human-animal interactions (HAIs) can bring about an array of benefits for the health and well-being of both humans and animals (Rault et al., 2020; Gee et al., 2021). However, the underlying mechanism of what brings about these benefits remains poorly understood, possibly due to the lack of available methodological approaches to ascertain the positive valence of interactions. Therefore, complementary approaches using both behavioural and physiological indicators are needed to investigate the distinguishing characteristics of positive interactions. Studying the neurochemical basis of social interactions can help to elucidate their proximate mechanisms (Carter and Keverne, 2002). In this study, we focused on neurotransmitters that are known to have key roles in positive social interactions: oxytocin, β-endorphin, dopamine, and serotonin.

Oxytocin (OT) is a nonapeptide that acts as a neurotransmitter in the central nervous system (CNS) and as a neurohypophysial hormone in the peripheral nervous system (PNS) (MacDonald and MacDonald, 2010). It has received considerable attention in recent decades for its central effects on the modulation of social behaviour (Menon and Neumann, 2023). Oxytocin can have anxiolytic effects but can also trigger threat-orienting responses and be associated with increased anxiety (Shamay-Tsoory and Abu-Akel, 2016). It has been posited that the apparent contradictions in OT’s effects is due to it having an overarching role in regulating the social salience of cues, whereby its effects are context-dependent (e.g. cooperation vs. competition, familiar vs. unfamiliar) and can be modulated by individual differences (Shamay-Tsoory and Abu-Akel, 2016). In the realm of social behaviour, OT is involved in the circuitries that modulate social preference, social reward, and social memory (Menon and Neumann, 2023). For instance, gentle physical interactions with a familiar human increased OT concentration in dogs (Rehn et al., 2014) and pigs (Rault, 2016). The majority of studies on the neuroendocrine basis of sociality have focused on OT, and some suggest that this overemphasis overlooks the potential confounding effects of other neurotransmitters that also play a role in social relationships (Pearce et al., 2017), such as opioids (Machin and Dunbar, 2011).

Endogenous opioids are a family of neuropeptides best known for their primary role as neurotransmitters in the CNS modulating pleasure and pain (Leknes and Tracey, 2008). A particular class of endogenous opioids, β-endorphins (β-END), have a high affinity to μ-opioid receptors that are present in high density in the hedonic hotspot of the brain (Peciña, 2008; Fields and Margolis, 2015). Thus, β-END are responsible for the hedonic “liking” of consummatory rewards, eliciting the feeling of pleasure (Machin and Dunbar, 2011), but also have a role in the “wanting” of rewards (Peciña, 2008). This extends to social rewards, where β-END concentrations have been shown to increase following allogrooming in monkeys that have previously been socially isolated (Keverne et al., 1989). Given that opioids do not readily cross the blood-brain-barrier (Machin and Dunbar, 2011), they need to be sampled centrally, and therefore have been difficult to measure directly. Consequently, little is known about their endogenous release patterns or function in regards to sociality, compared to other neurotransmitters that can be sampled peripherally (e.g. OT in saliva (Jong et al., 2015; Lürzel et al., 2020; MacLean et al., 2018)).

Dopamine (DA) is a catecholamine predominantly known for its role as a neurotransmitter in the CNS regulating movement, motivation, and reward (Juárez Olguín et al., 2016; Klein et al., 2019). Dopamine influences social behaviour through its key role in mediating reward in the mesolimbic system (Juárez Olguín et al., 2016; Klein et al., 2019), although its exact function continues to be heavily debated (Wise, 2004; Berridge and Kringelbach, 2008; Lerner et al., 2021; Gershman et al., 2024). Nevertheless, it is generally agreed that DA exerts its influence on key reward components, motivation and reinforcement learning, through mechanisms of reward prediction error signalling (*i.e.* whether the reward is better or worse than expected). Dopamine ramps (*i.e.* increases as the subject increases proximity to the reward) can be observed as value signalling of the expected reward (Berke, 2018; Gershman et al., 2024). These changes in DA are accompanied by covarying behavioural proxies of motivation such as the latency to respond to cues (Berke, 2018; Klein et al., 2019; Gershman et al., 2024).

Serotonin (5-HT) is a monoamine that acts as a neurotransmitter in the CNS and as a hormone in the PNS (Kanova and Kohout, 2021). Despite most 5-HT receptors being located in the enteric nervous system (Roberts et al., 2020), 5-HT is best known for its effect on neuropsychology including mood, reward, and attention amongst others (Berger et al., 2009; Kanova and Kohout, 2021). Indeed, depletion of 5-HT in pigs led to pessimistic-like behaviour in a cognitive bias paradigm, which highlights its role in cognitive-emotional processing (Stracke et al., 2017). Serotonin is a key player in social interactions through its influence on social adaptation (Duerler et al., 2022) and social reward (Kranz et al., 2010). For instance, optogenetic stimulation of dorsal raphe 5-HT neurons projecting to the nucleus accumbens increases mice sociability (Walsh et al., 2018). However, studies have often found contradicting results regarding serotonin’s role in reward processing, alluding to its multidimensionality in modulating responses to stimuli (Liu et al., 2020).

Although often experimentally studied in isolation, each neurotransmitter system seldom acts alone but rather in an interactive, and possibly synergistic, way. The oxytocinergic system interacts with the opioidergic system to regulate social behaviour (Putnam and Chang, 2022). Oxytocin also activates dopaminergic and serotonergic pathways, reinforcing social interactions (Shamay-Tsoory and Abu-Akel, 2016; Menon and Neumann, 2023). For example, co-activation of DA and OT are necessary for pair bond formation in female prairie voles (Liu and Wang, 2003; Loth and Donaldson, 2021). Dopamine and 5-HT also show interactive effects in reinforcement (Cardozo Pinto et al., 2024) and reward (Fischer and Ullsperger, 2017). Therefore, it is important to consider these neurotransmitters simultaneously. To our knowledge, no studies have investigated endogenous changes in these four neurotransmitters simultaneously within the context of social interactions.

Most neurotransmitters remain challenging to measure non-invasively given that their peripheral concentrations may not reflect changes at the central level (OT: (Veening et al., 2010; Jurek and Neumann, 2018; Menon and Neumann, 2023); opioid: (Machin and Dunbar, 2011; Veening et al., 2010); DA: (Elsworth et al., 1987); 5-HT: (Egri et al., 2020)). Cerebrospinal fluid (CSF) measures have been considered to reflect circulating brain neurotransmitter concentrations (OT: (Veening et al., 2010); opioid: (Dunbar, 2010); DA: (Ruckebusch and Sutra, 1984); 5-HT: (Moncrieff et al., 2023)). Sampling via a spinal catheter, for example, is a feasible and minimally invasive method to sample CSF repeatedly in conscious and freely moving animals. This has already been done successfully in pigs (Rault, 2016; Bergadano et al., 2019), a species of increasing popularity in neuroscience research for its anatomical similarities to the human brain (Lind et al., 2007).

Studies investigating the changes in the proteome are quite recent and allow moving beyond the traditional targeted analysis of a few neurochemicals, and particularly beyond the analysis of a few favourite molecules that may hinder our comprehensive understanding of neuroendocrine processes, illustrated by the overemphasis on oxytocin in the literature on social interactions. Quantitative proteomics analysis enables screening protein changes may reveal novel proteins linked to positive interactions. A study found changes in neurotransmitter proteins in pigs due to a putatively positive intervention (environmental enrichment) (Arroyo et al., 2016), but the effect of HAIs on the proteome has not yet been explored.

In this study, we aimed to determine the specific changes in neurotransmitter (OT, β-END, DA and 5-HT) concentrations in relation to positive HAIs using the pig as a model. In a previous study (Rault, 2016), positive human-pig interactions with a familiar human were shown to increase CSF OT, but the effects of familiarity and valence of the interaction were confounded, and OT is known to be sensitive to familiarity vs. novelty (e.g. (Tops et al., 2013b)). Therefore, in this study we used a 2 × 2 factorial design to disentangle the effects of human familiarity (familiar (‘F’) vs. unfamiliar (‘U’)) and type of interaction offered to the pig (human provides positive contact (‘p’) vs. human ignores the pig (‘i’)). We hypothesised that OT underlies social selectivity, opioid (β-END) and 5-HT underly the hedonic property of positive interactions, and DA underlies the motivation to interact. We expected OT to increase following interactions with a familiar human (‘F’ conditions) compared to unfamiliar human (‘U’ conditions), DA to increase in all conditions but more so with positive contacts (‘p’ conditions), and β-END and 5-HT to increase during positive contacts (‘p’ conditions) compared to neutral contacts (‘i’ conditions). No specific predictions were made for the changes in proteome, as this part of the study was explorative.

## 2. Material and Methods

### 2.1. Ethical approval

This experiment was conducted in accordance with the national legislation and the European Directive 2010/63/EU. It was approved by the BMBWF (Bundesministerium für Bildung, Wissenschaft und Forschung) under the approval number 2021-0.530.305, and by the ethical committee of the University of Veterinary Medicine Vienna under the approval number ETK 073/05/2021.

### 2.2. Animals, housing and feeding

A total of 36 female pigs (*Sus Scrofa domesticus;* Swiss Large White × Pietrain breed) was used in three batches of 12 pigs. The pigs were recruited at weaning (4 weeks of age) and housed at the Swine Clinic at the main campus of Vetmeduni. The pigs were selected based on good health condition and avoiding weight extremes.

Pigs were housed in individual home pens (1.10 × 1.74 m) until the end of the experiment to avoid manipulation of the catheter by another pig. Partitions between pens allowed visual and olfactory access as well as snout to snout contact with neighbouring pigs to minimise the stress of single housing. Water and feed (*i.e.* standard weaner diet; 17.5% crude protein, 7% crude fat; Garant-Tiernahrung GmbH, Austria) were provided *ad libitum*. Straw bedding on the full concrete floor was renewed daily to ensure a clean and comfortable environment, and extra enrichment was provided in the form of a braided jute rope (1 m long) hung from the pen door, an orange dog toy ball (Airflow ball, Dog Crest, 7.6 cm diameter) and a rotation of toilet paper rolls filled with hay or banana chips delivered daily. Additional gentle contacts were provided by the humans to the pigs at the end of each day in order to decrease the potential stress of single housing. The windows allowed natural lighting of the room and the pens, and room temperature was maintained at around 21°C by heaters with heat lamps suspended about 60 cm from the ground.

### 2.3. Experimental design

#### 2.3.1. Habituation to human contacts and to the test pen

Starting from 3 days post-weaning (*i.e.* 5 weeks of age), pigs were gradually habituated to human presence, human contact, and to the test pen over 10-minute sessions conducted twice daily over two weeks.

The two experimenters (*i.e.* the handler and the familiar human, both women) were involved in the sessions of habituation to human presence and human contacts, but only the familiar human provided contacts in the test area. During the habituation sessions, the experimenter entered the home pen, kneeled in the corner opposite to the door and talked softly to the pigs. If the pig initiated contact with her, she presented her hand to it and then gently stroked and/or scratched it (first on the head area and slowly moved towards the rear of the body, including belly). These contacts included belly-rubbing if the pig rolled on its back and exposes its belly. Touching the ears of the pigs was avoided as it may not be perceived as a pleasant contact (personal observations). Sudden movements and loud voices were avoided at all times. However, if pigs started to bite the human, the human experimenter withdrew from the pig, and gently tapped the insisting pig on its snout to discourage it from biting.

After three sessions of habituation to human contact in their home pen, each pig was gently guided, without force, individually by the handler to the test area (located in the same room as the home pens) where the familiar human entered after the pig and followed the same procedure as in the home pen.

During habituation, it was planned to terminate the session earlier if the pig showed signs of distress (*i.e.* three consecutive acts of defecation and/or urination, five high-pitched vocalisations within 1 min, or two attempts to escape). However, this did not happen with any of the pigs.

### 2.4. Surgical procedures for spinal catheter placement

#### 2.4.1. Sedation and anaesthesia

Pigs underwent the spinal catheter placement under anaesthesia performed by a certified veterinary anaesthesiologist (Dr. Attilio Rocchi). Anaesthesia followed a 3-stage strategy designed to minimize anaesthesia load, provide precise individualized modulation of anaesthetic depth, and prevent harm and suffering in the event of unexpected issues (*e.g.* respiratory problems). Pigs received ketamine hydrochloride (Narketan®, 7 mg/kg Body Weight [BW]) and Azaperon (Stresnil®, 1-2 mg/kg BW) intramuscularly in the neck region behind the ear (brachiocephalic muscle) to achieve deep sedation. Subsequently, a 22G venous catheter was placed in the external auricular vein of the pig and secured with tape. Anaesthesia was then induced with ketamine hydrochloride (Narketan®, 5 mg/kg BW) and Midazolam (2 mg/kg BW) administered intravenously. Induction was followed by tracheal intubation with a cuffed endotracheal tube and delivery of isoflurane in 50% oxygen through a circular system for controlled maintenance of anaesthesia. Under general anaesthesia, a 22G venous catheter was placed in the central auricular artery for invasive blood pressure monitoring. To ensure that pigs were fully anesthetised, the anaesthetist monitored the loss of the righting reflex and loss of the palpebral eye reflex by touching the side of the eye before proceeding. Eye lubrication was administered to the eyes to prevent drying of the cornea during the surgery. In the case of apnoea (cessation of breathing for 30 sec) and/or hypoventilation (End-Tidal CO2 (EtCO2) > 50 mmHg), intermittent positive ventilation was started to maintain EtCO2 between 40-45 mmHg. Heart rate, electrocardiogram, peripheral oxygen saturation, respiration rate, EtCO2, blood pressure, and body temperature were monitored with a multi-parameter monitor (HP CMS-2000 Anesthesia monitor) throughout the surgery. Intraoperative analgesia was provided with meloxicam (Metacam®, 0.4 mg/kg BW), administered intravenously after placement of the venous catheter, and local infiltration of Lignocaine (3 mL of Xylanaest purim 2%, Gebro Pharma GmbH) prior to placing the spinal catheter.

#### 2.4.2. Placing of the spinal needle and catheter

The placement of the spinal catheter consisted of making a small incision in the skin of 0.5 cm with a scalpel blade to ease the crossing of the spinal needle through the pig’s tough skin. A spinal needle (B-Braun® SPINOCAN Spinal needle, 16 Ga × 8.9 cm, B-Braun Medical, Boulogne Billancourt, France) was inserted into the spinal subarachnoid space by lumbar puncture through the lumbar interspace, the needle advancing until piercing the dura mater. Placement was verified by dripping of CSF through positive pressure before fitting a spinal catheter (B-Braun® PERIFIX Epidural catheter set, 18 Ga × 100 cm, B-Braun Medical, Boulogne Billancourt, France) 2 to 5 cm inside the subarachnoid space, which was secured using a tape externally glued to the outer edge of the skin. The tip of the catheter was secured by a tape sutured to the skin to ensure that the catheters did not slip out, and the external part of the catheter was kept in a small pouch, specifically designed for this experiment and glued on the back of the pig, to ensure the catheter was kept clean at all times.

#### 2.4.3. Recovery after placement of the spinal catheter

Each pig was monitored every 10 min until they were able to stand, walk, and eat treats from the experimenter’s hand. They continued to be monitored every 30 min for 4 hours following the surgery.

After the catheter placement procedure, the pigs were given two days to recover, during which they received daily positive human contacts in their home pen. In order to monitor signs of pain after spinal catheter placement and administer pain-relief treatment if needed, the pain assessment scale for swine developed by Swindle (Swindle, 2009) was used with an assessment done daily at the time of flushing the catheters. No pain reaction (i.e. to tactile check around the insertion point) was associated with the placement of the catheter. However, two pigs reacted (i.e. made a sudden movement) during the flushing of the catheter: one of them pulled the catheter out and the other one had a non-functional catheter, in both cases we stopped trying to sample or flush for the rest of the trial. Two pigs experienced temporary paralysis of the back legs (i.e. for 4 and 7 days) and one pig did not recover the use of its back legs; those pigs were excluded from the trial and not subjected to any test or sampling procedures, and instead received anti-inflammatory drugs as the veterinarian suspected that this was caused by an inflammatory reaction around the nerves surrounding the spinal cord.

The handler flushed the catheters daily throughout the experiment with a 0.9% sterile saline solution. Pigs were not restrained or handled during the flushing procedure, which lasted between 2 and 5 minutes and consisted of 1) wiping the connector with an alcohol swab, 2) drawing liquid from the catheter using a syringe and discarding at least 0.20 ml of liquid (corresponding to the catheter dead space), 3) switching syringe and collecting 0.25ml of CSF, which was then stored in an Eppendorf tube at −80°C, and 4) injecting 0.20 ml of sterile saline solution to fill the catheter dead space.

After two post-surgery recovery days, each pig with a functional catheter (*i.e.* from which CSF could be successfully sampled as assessed during daily flushing of the catheter) and showing no signs of pain or abnormal behaviour (e.g. altered gait, lethargy indicative of post-operative effects) was subjected to testing.

### 2.5. Testing

#### 2.5.1. Procedure

Each pig was individually subjected to four different 10-minute tests according to a within-subject, 2 × 2 factorial design to disentangle the effects of human familiarity (familiar or unfamiliar human) and type of interaction offered to the pig (positive contacts or ignoring). On each testing day, the pigs were subjected to testing in the morning in a fixed order in which they were randomly assigned one of the following test situations:

“Familiar positive” (Fp): A familiar human, who has previously interacted with the pig during habituation, provides positive contacts;

“Unfamiliar positive” (Up): An unfamiliar human who also delivers positive contacts;

“Familiar ignore” (Fi): A familiar human who is unresponsive to the pig (control 1);

“Unfamiliar ignore” (Ui): An unfamiliar human who is unresponsive to the pig (control 2).

In all test situations, the pig was moved individually into the test pen first and the human entered a few seconds later. The test began the moment the human walked to the corner opposite the pen door. The human then sat down and remained in the corner for the duration of the test session, allowing the pigs to voluntarily approach.

In both tests where the human provides positive contacts (Fp and Up), the pig was encouraged to approach through vocal (soft talking voice) and physical solicitation cues (tapping of the fingers on tights or floor, small hand and arm gestures avoiding big and fast movements). The pig was given gentle tactile contact (stroking, rubbing, and scratching) when within an arm’s reach. The unfamiliar humans were instructed by the familiar human about how to interact with the pigs, to avoid major differences in interaction style and make sure the difference in the pigs’ response was more due to differences in human familiarity than to differences in the way humans interacted with the pig.

In both tests where the human was ignoring the pig (Fi and Ui), the human remained seated with their arms folded and hands tucked under their armpits to prevent manipulation of the hand by the pig. The human maintained a soft gaze to the wall on the opposite side of the pen and avoided eye contact with the pig.

#### 2.5.2. CSF sample collection

A maximum of 1 ml of CSF was sampled by trained personnel (OS) before (65 min [B1] and 5 min [B2] pre-test) and after the start of the test (10 [T10], 30 [T30], 60 [T60], 120 [T120], 240 [T240] min post-test) (Figure 1), as in (Rault, 2016), using a graduated syringe to allow estimating the quantity withdrawn.

**Figure 1.**
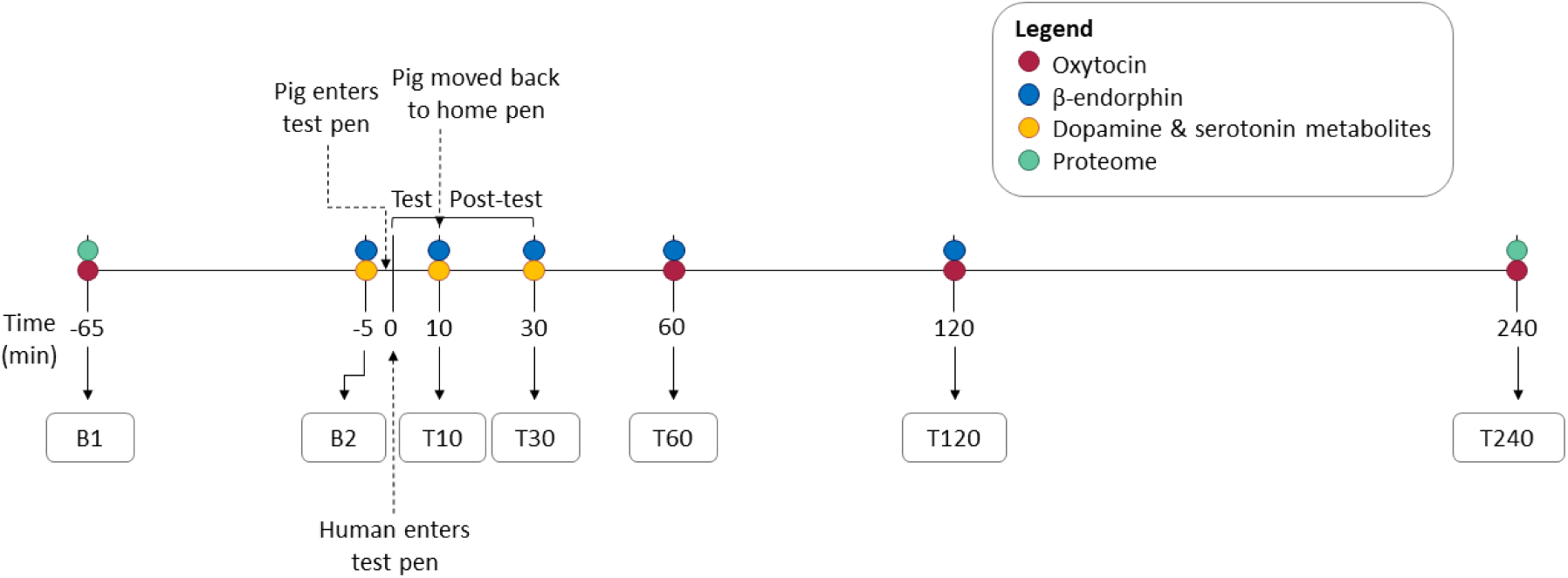
Test session timeline showing the seven cerebrospinal fluid (CSF) sampling time points. The 10-min test began at time 0. Samples were collected at two timepoints before the test (B1 at 65 minutes pre-test and B2 at 5 minutes pre-test), and five timepoints post-test (T10, T30, T60, T120, T240 minutes). Given the limited volume of CSF that could be collected at each timepoint, and the volumes required for each neurotransmitter analyses, each sample was allocated to specific analyses based on the expected time dynamics of change in each neurotransmitter: OT (B1, T60, T120, T240), β-END (B2, T10, T30, T60, T120), DA and 5-HT metabolites (B2, T10, T30), Proteome (B1, T240). B = baseline; T = time.

These CSF samples were analysed for their concentrations of OT, β-END, DA and 5-HT metabolite concentrations and for proteomics analysis. The CSF sampling procedure was similar to the flushing procedure, except that 1 ml of CSF was withdrawn, placed in an Eppendorf tube on dry ice, and stored in the −80°C freezer until analysis. Samples destined to proteomics analysis contained protease inhibitor (cOmplete™ Tablets, Mini EDTA-free, EASYpack; Roche Diagnostics GmbH, Germany) to prevent the degradation of proteins.

### 2.6. Data analysis

#### 2.6.1. CSF sample analysis

Samples were analysed for OT concentrations by ELISA (Cayman Chemical Oxytocin ELISA kit No 500440). Each sample was analysed in duplicates, and samples from the same pig were analysed within the same assay run in order to avoid variation due to the inter-assay CV and given the within-subject experimental design. The intra-assay CV was 4.43% on average [range: 0.09 - 13.90%].

Samples were analysed for β-endorphin concentrations by EIA (Phoenix Pharmaceuticals Endorphin beta (Porcine) EIA Kit EK-022-032). Each sample was analysed in duplicates, and samples from the same pig were analysed within the same assay run in order to avoid variation due to the inter-assay CV and given the within-subject experimental design. The intra-assay CV was 4.26% on average [range: 0.04 – 18.99%]. Some extra samples were pooled and run across plates to calculate the inter-assay CV, which was 27.61% on average [range: 0.65 – 56.14 %].

Concentrations of DA’s metabolites 3,4-dihydroxyphenylacetic acid (DOPAC) and homovanillic acid (HVA), and the 5-HT’s metabolite 5-hydroxyindole-3-acetic acid (5-HIAA) were determined in duplicate by high-performance liquid chromatography (HPLC) with electrochemical detection. For deproteinisation, 100 µl CSF were shortly vortexed with 2.5 µl 11.7 M perchloric acid followed by centrifugation at 37,000 *g* for 10 min at 4 °C. From the supernatants, aliquots of 50 µl were then injected directly into the HPLC system. The HPLC system (LC-10 series, SHIMADZU, Duisburg, Germany) was equipped with a 125 x 4 mm reversed phase column packed with Prontosil C18 AQ (Bischoff Analysentechnik, Leonberg, Germany). Separation was achieved with a mobile phase consisting of 58 mM sodium hydrogen phosphate buffer containing 1.2 mM octanesulfonic acid, 0.3 mM EDTA, 0.2 mM potassium chloride, and 9% methanol at pH 2.6. Column temperature was 40 °C and a flow rate of 1.2 ml/min was set. Electrochemical detection was performed using a DECADE II detector with a SenCell (Antec Scientific, Alphen, The Netherlands) at a potential of 600 mV. A two-point calibration with external standards was used. In the CSF samples, DA and 5-HT were not detectable and the metabolites DOPAC, HVA and 5-HIAA were used as proxy measures of dopaminergic and serotonergic activity. The intra- and interassay coefficients of variation (CV) were 1.9 % and 11.6 % for DOPAC, 3.3 % and 5.6 % for HVA, and 0.4 % and 3.2 % for 5-HIAA, respectively.

##### Proteomics

Sample preparation and analysis was conducted at the Proteomics and Metabolomics Facility of the Research Center for Molecular Medicine of the Austrian Academy of Sciences (CeMM). Discovery proteomics analysis (label free) consisted of comparing 4 pigs across 4 time points in a label free data-independent analysis (DIA) liquid chromatography-mass spectrometry (LC-MS) approach to identify significant changes in protein abundance.

Protein amount in cerebrospinal fluid (CSF) was quantified using Pierce BCA Protein Assay (ThermoFisher Scientific, San Jose, CA). For this, 15 µg of protein were used for SP3 digest. Reduction was performed with DTT (1h, 56°C) and alkylation with IAA (30 min, RT). Peptides were cleaned up with Pierce Peptide Desalting Columns (ThermoFisher Scientific, San Jose, CA) and resuspended to 100 ng/µl, i.e. 500 ng were injected for each sample (5 µl).

Mass spectrometry analysis was performed on an Orbitrap Fusion Lumos Tribrid mass spectrometer (ThermoFisher Scientific, San Jose, CA) coupled to a Dionex Ultimate 3000 RSLCnano system (ThermoFisher Scientific, San Jose, CA) via a Nanospray Flex Ion Source (ThermoFisher Scientific, San Jose, CA) interface. Peptides were loaded onto a trap column (PepMap 100 C18, 5 μm, 5 × 0.3 mm, ThermoFisher Scientific, San Jose, CA) at a flow rate of 10 μL/min using 0.1% TFA as loading buffer. After loading, the trap column was switched in-line with a C18 analytical column (2.0 µm particle size, 75µm IDx500mm, ThermoFisher Scientific, San Jose, CA). The column temperature was maintained at 50 °C using a PRSO-V2 column oven (Sonation, Biberach, Germany). Mobile phase A consisted of 0.4% formic acid in water, and mobile phase B consisted of 0.4% formic acid in a mixture of 90% acetonitrile and 10% water.

Separation was achieved using a four-step gradient over 91 minutes at a flow rate of 230 nL/min (increase of initial gradient from 4% to 24% solvent B within 82 minutes, 24% to 36% solvent B within 8 minutes, 36% to 100% solvent B within 1 minute, and 100% solvent B for 6 minutes before equilibrating to 4% solvent B for 18 minutes before the next injection).

In the liquid junction setup, electrospray ionization was enabled by applying a voltage of 1.8 kV directly to the liquid being sprayed, and a non-coated silica emitter was used.

The mass spectrometer was operated in a data-independent acquisition mode (DIA). For MS2 acquisition, we collected a 350–1650 m/z survey scan in the Orbitrap at 120 000 resolution (FTMS1), the AGC target was set to 200% and a maximum injection time (IT) of 100 ms was applied. Precursor mass range for data-independent analysis was set to 380-880 m/z. Precursor isolation was performed with quadrupole. Isolation window size was set to 5 m/z with an overlap of 2 m/z. The RF lens was set to 30%. MS2 was measured in the Orbitrap with a resolution of 30 000.

High-energy collision-induced dissociation (HCD) fragmentation technique was used at a stepped collision energy of 24, 28 and 30%. The normalized AGC target was set to 200% with a dynamic maximum IT with at least 10 points across the peak. Xcalibur Version 4.3.73.11 and Tune 3.4.3072.18 were used to operate the instrument.

The data were analysed using PEAKS 11.0 (Studio version, build 20230414) (Bioinformatics Solutions Inc., Waterloo, Canada) in order to calculate the normalised protein abundance value for each identified protein within the data-independent analysis label free quantification dataset (DIA). The protein databases used for the searches included a *Sus scrofa* database downloaded from Uniprot (www.uniprot.org) in September 2023 (taxonomy ID 9823) as well as common contaminant (cRAP) database (downloaded from https://www.thegpm.org/crap/). Following parameters were applied: enzyme trypsin, maximum of two missed cleavage sites, digest mode specific, precursor mass tolerance 10 ppm and fragment mass tolerance 0.02 Da, variable modification: oxidation (+15.995 Da on methionine), fixed modification: carbamidomethyl (+57.021 Da on cysteine).

#### 2.6.2. Behavioural observations

Video recording was done using a handheld camcorder (Sony CX-730) fixed on a tripod above the test pen for recording of the test sessions. The videos were then analysed by a single observer and coded for behaviour using the software BORIS v 8.11.2 (Friard and Gamba, 2016) (see full ethogram in Supplementary Material, Table S1). The Cohen’s Kappa score for the intra-observer reliability was 0.83 on average [range: 0.61-0.96]. An external observer also coded some videos to establish the inter-observer reliability, Cohen’s Kappa score was 0.86 on average for the behaviours of interest [range: 0.69–0.96]. Only the contacts initiated by the pig were used for the statistical analysis, but the complete ethogram used for the video analysis can be found in the Supplementary materials (Table S1). “Contact by pig” was defined as the pig making contact with the human using its snout (sniffing, nosing, oral manipulation), or the pig was either standing with its body against the human or lying/sitting on the human (i.e. body contact), or placing at least its front legs on the human (i.e. climbing on human).

### 2.7. Statistical analysis

#### 2.7.1. Behaviour and neurotransmitters

A coding system was used to anonymize the samples and datasets, which allowed blinding all personnel involved in the laboratory and statistical analyses to the treatments.

All data were analysed using mixed models with the fixed effects of human familiarity, contact type, time of sampling (‘time’), the three-way interaction of human familiarity × contact type × time of sampling, and batch. The models also included the random intercept of pig and the random slopes of human familiarity, time and contact type.

We used a full-null model comparison (based on a likelihood ratio test; (Dobson and Barnett, 2018)) to test the effects of human, contact type, their interaction, and their respective interactions with time (Forstmeier and Schielzeth, 2011), and avoid cryptic multiple testing. Only the random effect of pig and the random slopes of time and contact type were included in the null model. Satterthwaite approximation was used to test the effect of individual fixed effects (Luke, 2017) with the function lmer of the package lmerTest (version3.1-3, (Kuznetsova et al., 2017)) and a model fitted with restricted maximum likelihood.

Prior to fitting the model, the quantitative predictors were inspected for whether their distributions were roughly symmetrically distributed. The distribution was left-skewed for most variables, but given that data transformation (log, square root) did not improve their distribution, or the fitness of the statistical models, these variables were left untransformed.

After fitting the model, the assumptions of normally distributed and homogeneous residuals were visually inspected on a QQ-plot (Field, 2005) of residuals and residuals plotted against fitted values (Quinn and Keough, 2002). No deviations from these assumptions were observed. Collinearity, determined for a model lacking the interaction, appeared to not be an issue (maximum Variance Inflation Factor: 1.59; (Quinn and Keough, 2002)).

Model stability was assessed on the level of the estimated coefficients and standard deviations by excluding the levels of the grouping factors one at a time, using a function written by Rodger Mundry. This revealed the models to be of acceptable stability for all models.

We fitted the model in R (version 4.1.2; (R Core Team, 2024)) using the function lmer of the package lme4 (version 1.1-27.1; (Bates D., Maechler M., Bolker B., 2015)). We determined variance inflation factors using the function *vif* in the package *car* (version 3.0-11; (Fox et al., 2012)).

The sample for each model encompassed 109, 154 or 163 values taken from 10 pigs out of three batches on three (HVA, HIAA, DOPAC), four (OT) or five (β-END) timepoints, respectively.

Due to the explorative nature of our work, no correction for multiple comparisons was applied. Significance was considered when the P-value was smaller than 0.05 and a tendency was considered when the P-value was between 0.05 and 0.1.

#### 2.7.2. Proteomics

All statistical analyses were performed in R (v4.2.3. (R Core Team, 2024)).

Data were prepared using functions from packages *tidyr* v1.3.0 (Wickham et al., 2023b), *dplyr* v1.1.4, *reshape2* v1.4.4 (Wickham, 2007) and *stringr* v1.5.1r_1.5.1 (Wickham, 2023).

Figures were produced with base R and functionality from package *ggplot2* v3.4.4 (Wickham, 2016) and exported in scalable vector graphics format via package *svglite* v2.1.3 (Wickham et al., 2023a). We used colourblind-friendly palettes from *RColorBrewer* v1.1-3 (Neuwirth, 2022).

We produced heatmaps of unsupervised clustering, based on Euclidean distances and correlation coefficients, applying different clustering methods, on 1619 row-wise z-transformed proteins with function *pheatmap* from package *pheatmap* 1.0.12 (Kolde, 2019).

We used the function *fitExtractVarPartModel* in package *variancePartition* v1.28.9 (Hoffman and Roussos, 2021) to simultaneously estimate the variances of the interaction of time (factor levels: pre-test measure B1 and post-test measure T240) with human familiarity (factor levels: familiar and unfamiliar) and contact type (ignore and positive), resulting in a total of 8 factor levels, 7 dummy coded centered uncorrelated random slopes of the interaction term, random intercept term of animal (10 factor levels), and as potential sources of technical noise housing batch (3 factor levels) and date of proteomics quantifications (12 factor levels). For each protein the amount of variance explained by each effect is estimated and visualized as violin plots across all 1619 genome wide proteins. We summarized the variances across all seven dummy coded centred random slopes, which represent interactions between our interaction of time, human familiarity and contact type and the random animal effects. Protein quantifications of 1619 proteins were log10 transformed after adding a constant of 1 to all observations.

Differential protein expression analyses with function dream in package *variancePartition v1.28.9* did not reveal significant differences at a global 5% FDR threshold after multiple testing correction (data not shown).

## 3. Results

The summary of all statistical results can be found in the Supplementary Material (Table S2).

### 3.1. Oxytocin

There was a significant effect of contact type (F_1, 25.6_ = 8.4; P = 0.008), with the OT concentration being overall higher in the “ignore” condition than in the “positive contacts” condition (18.9 ± 1.36 ng/mL vs 16.4 ± 1.43 ng/mL; t_25.6_ = 2.89; P = 0.008).

There was a tendency for the interaction human familiarity × contact type (F_1, 119.8_ = 3.19; P = 0.08; Figure 2), with pairwise comparisons revealing that the OT concentration was lowest in the Up condition and was the highest in the Ui condition (15.3 ± 1.56 pg/mL vs 19.4 ± 1.49 pg/mL; t_70.6_ = 3.31; P = 0.008), and OT concentration tending to be lower in the Up condition than in the Fi condition (15.3 ± 1.56 pg/mL vs 18.4 ± 1.50 pg/mL; t_71.9_ = 2.53; P = 0.06). The OT concentration in the Fp condition was intermediate (17.4 ± 1.56 pg/ml) and did not differ from the other conditions (0.3 < P < 0.9).

**Figure 2.**
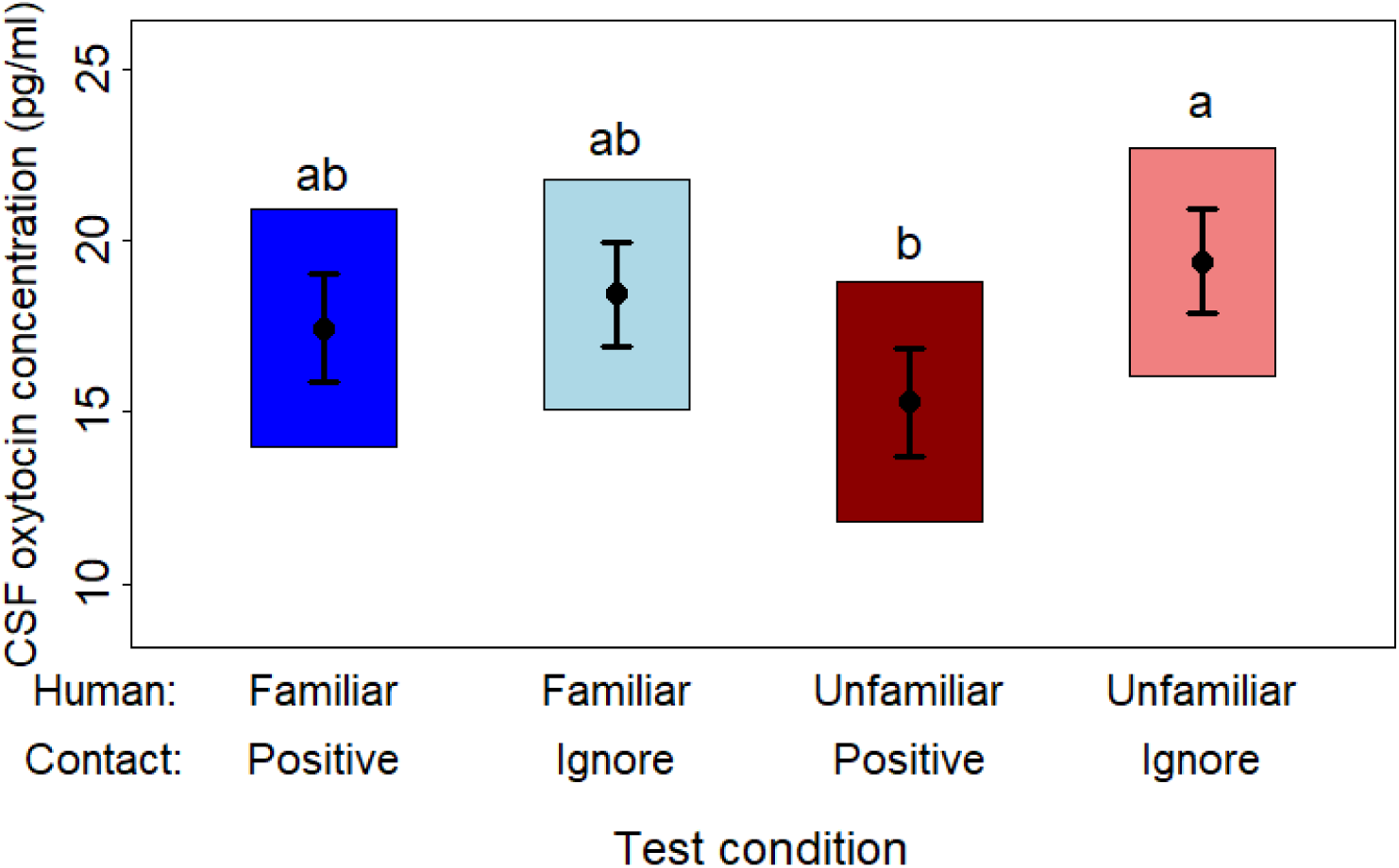
Overall effects of the test conditions (interaction between the human familiarity and the type of contacts) on the concentration of oxytocin. The graph shows the estimated means (dots) and standard errors (error bars), as well as the 95% confidence intervals (box) estimated from the statistical model. The statistical significance of the interaction between the two factors was P = 0.08. The pairwise significant differences (P < 0.05) are represented by different letters.

There was an overall significant effect of time (F_3, 25.4_ = 8.4; P < 0.001), with the OT concentration being lower 240 min post-test compared to 60 min (13.6 ± 1.07 vs. 19.8 ± 1.18; t_8.74_ = 5.059; P < 0.005) and 120 min (13.6 ± 1.07 vs. 19.5 ± 1.68 pg/mL; t_8.89_ = 4.012; P < 0.05) post-test, and tending to be lower 240 min post-test compared to pre-test (13.6 ± 1.07 vs. 17.3 ± 0.95 pg/mL; t_8.91_ = 3.015; P = 0.06). Finally, there was no effect of the human familiarity (F_1, 119.4_ = 0.5; P = 0.49), the interaction between contact type and time (F_1, 119.8_ = 1.07; P = 0.37), or the three way interaction human familiarity × contact type × time (F_1, 119.9_ = 1.34; P = 0.26; Figure S1).

### 3.2. 3,4-Dihydroxyphenylacetic acid (DOPAC) and homovanillic acid (HVA)

Analysis of the dopamine metabolite DOPAC revealed a tendency for the effect of the interaction between human familiarity and contact type (F_1,70.57_ = 3.31; P = 0.07, Figure 3), but pairwise comparisons were not significant. No other factors had a significant effect on DOPAC concentration: human familiarity (F_1,9.5_ = 0.42; P = 0.53), contact type (F_1,8.5_ = 0; P = 0.96), time (F_2,26.9_ = 0.23; P = 0.8), human familiarity × time (F_2,68.5_ = 0.58; P = 0.56), contact type × time (F_2,68.5_ = 0.86; P = 0.43), human familiarity × contact type × time (F_2,67.7_ = 0.02; P = 0.98; Figure S2).

**Figure 3.**
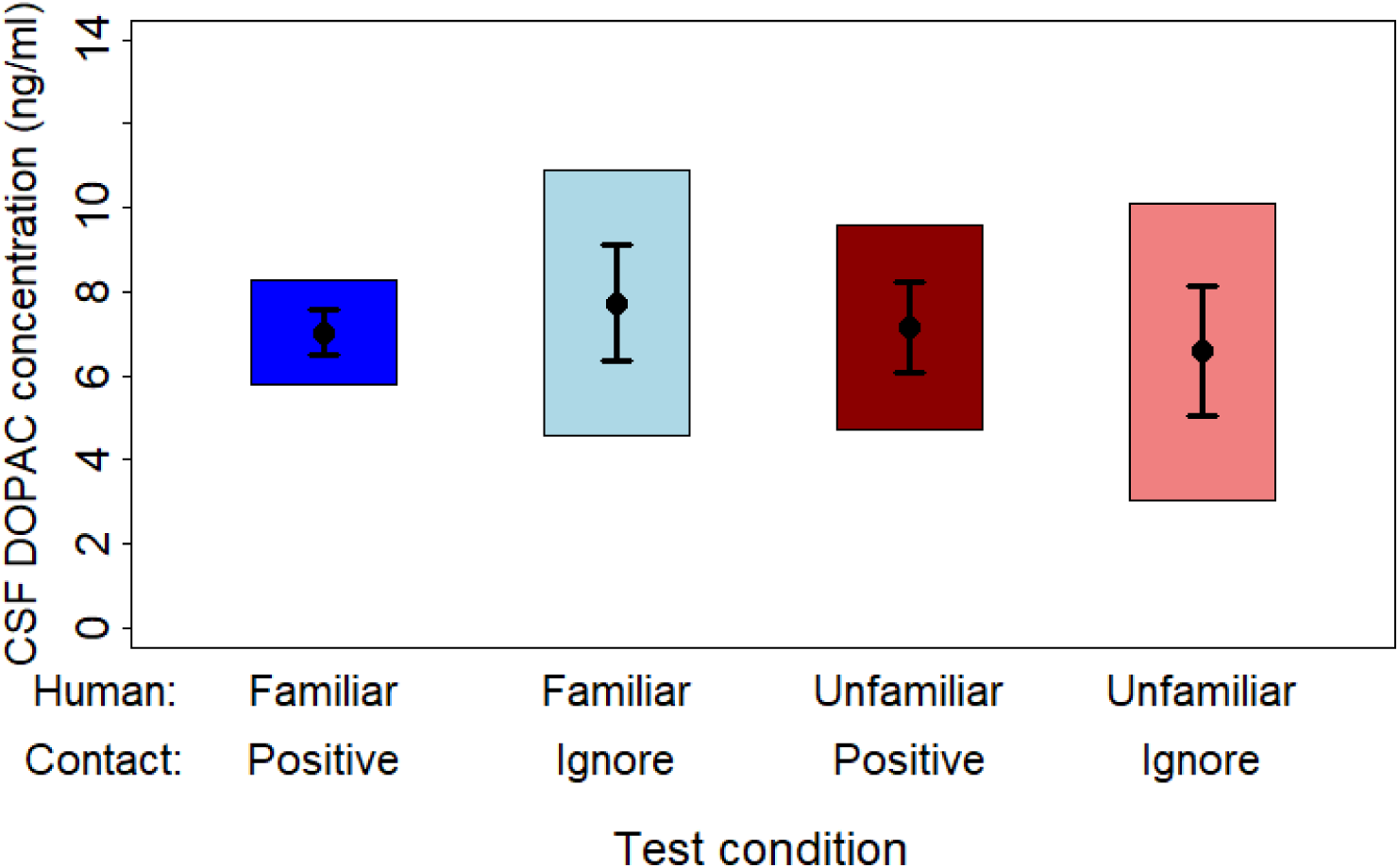
Overall effects of the test conditions (interaction between the human familiarity and the type of contacts) on the concentration of DOPAC. The graph shows the estimated means (dots) and standard errors (error bars), as well as the 95% confidence intervals (box) estimated from the statistical model. The statistical significance of the interaction between the two factors was P = 0.07.

For the dopamine metabolite HVA, there was a significant overall effect of time (F_2,18_ = 5.2; P = 0.017) where the concentration of HVA was higher 30 min post-test compared to pre-test (27.1 ± 3.38 ng/ml vs. 23.4 ± 3.32 ng/ml; t_8.32_ = 3.025; P < 0.05). The concentration of HVA 10 min post-test was intermediate (25.9 ± 2.84 ng/ml) and was not significantly different to the pre-test and 30 min post-test measures (P = 0.2 and P = 0.7, respectively). No other factors had a significant effect on the HVA concentration: human familiarity (F_1,9.1_ = 0.86; P = 0.38), contact type (F_1,8.9_ = 0.05; P = 0.83), human familiarity × contact type (F_1,71.6_ = 0.01; P = 0.92), human familiarity × time (F_2,69.8_ = 0.59; P = 0.56), contact type × time (F_2,70.6_ = 0.83; P = 0.44), human familiarity × contact type × time (F_2,70_ = 0.82; P = 0.44; Figure S3).

### 3.3. 5-Hydroxyindoleacetic acid (5-HIAA)

The analysis of the serotonin metabolite 5-HIAA concentration revealed a significant effect of the interaction between human familiarity and contact type (F_1,73.47_ = 5.02; P = 0.03, Figure 4), but pairwise comparisons did not show significant differences between the test conditions. No other factors had a significant effect on the 5-HIAA concentration: human familiarity (F_1,9.1_ = 0.58; P = 0.47), contact type (F_1,9_ = 0.07; P = 0.8), time (F_2,38.6_ = 2.13; P = 0.13), human familiarity × time (F_2,70.9_ = 0.23; P = 0.79), contact type × time (F_2,70.9_ = 0.01; P = 0.99), human familiarity × contact type × time (F_2,70.3_ = 0.31; P = 0.73; Figure S4).

**Figure 4.**
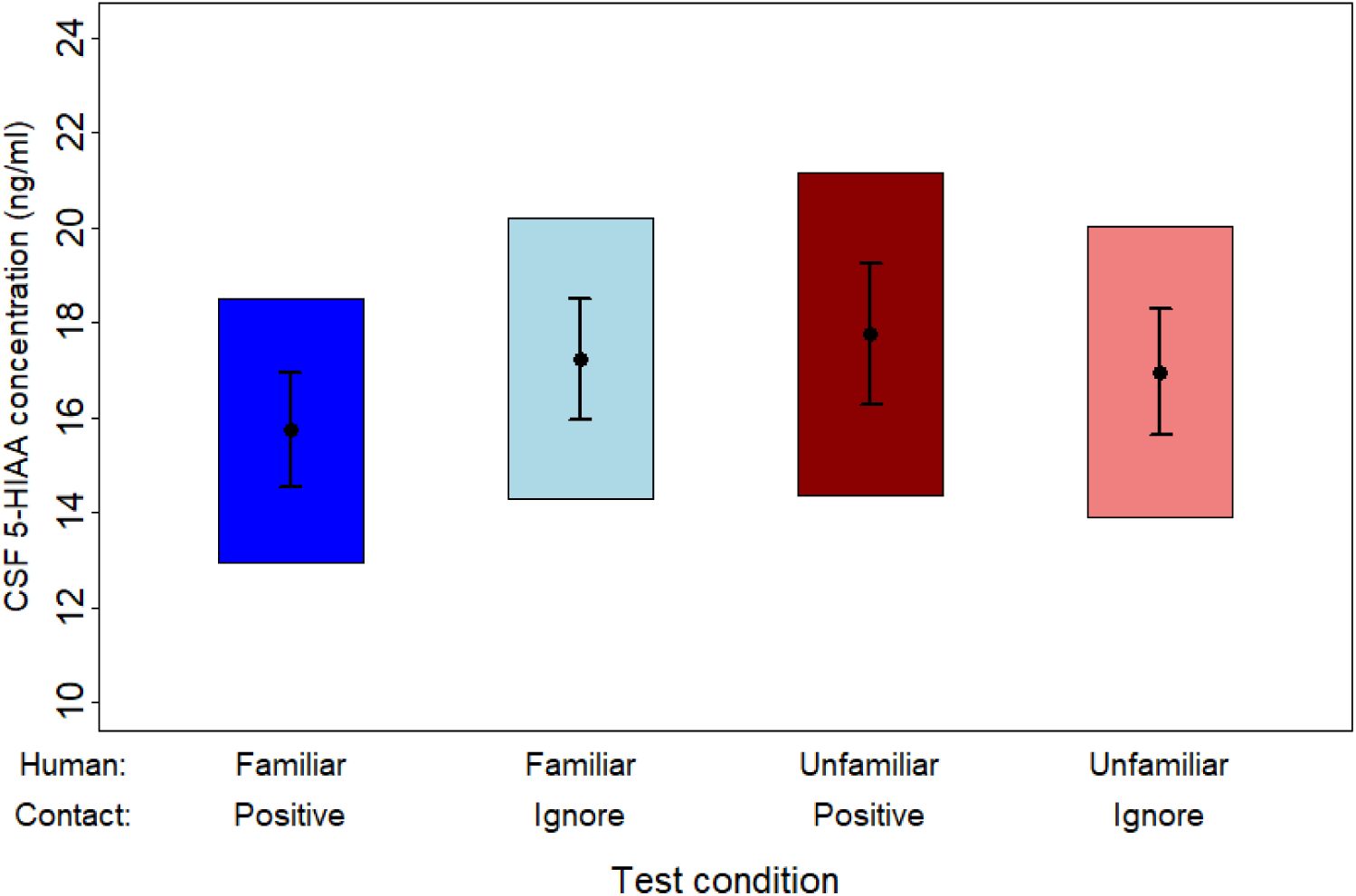
Overall effects of the test conditions (interaction between the human familiarity and the type of contacts) on the concentration of 5-HIAA. The graph shows the estimated means (dots) and standard errors (error bars), as well as the 95% confidence intervals (box) estimated from the statistical model. The statistical significance of the interaction between the two factors was P = 0.03 but pairwise comparisons were not significant.

### 3.4. *β*-endorphin (β-END)

There was a significant effect of the interaction between human familiarity × contact type on β-END concentration (F_1,118.59_ = 4.45; P = 0.04), but pairwise comparisons showed no significant differences between test conditions.

There was also a significant effect of the interaction human familiarity × time (F_4,116.1_ = 4.67; P = 0.001, Figure 5), as β-END concentration tended to be higher 1 h after interacting with the “familiar human” than with the “unfamiliar human” (0.16 ± 0.014 ng/ml vs. 0.10 ± 0.014 ng/ml; t_37.9_ = 3.3; P = 0.055). No other factors had a significant effect on β-END concentration: human familiarity (F_1,9.5_ = 1.53; P = 0.25), contact type (F_4,22.1_ = 0.35; P = 0.84), time (F_1,9.9_ = 0.61; P = 0.45), contact type × time (F_4,115.4_ = 0.36; P = 0.83), human familiarity × contact type × time (F_4,115.2_ = 0.69; P = 0.6; Figure S5).

**Figure 5.**
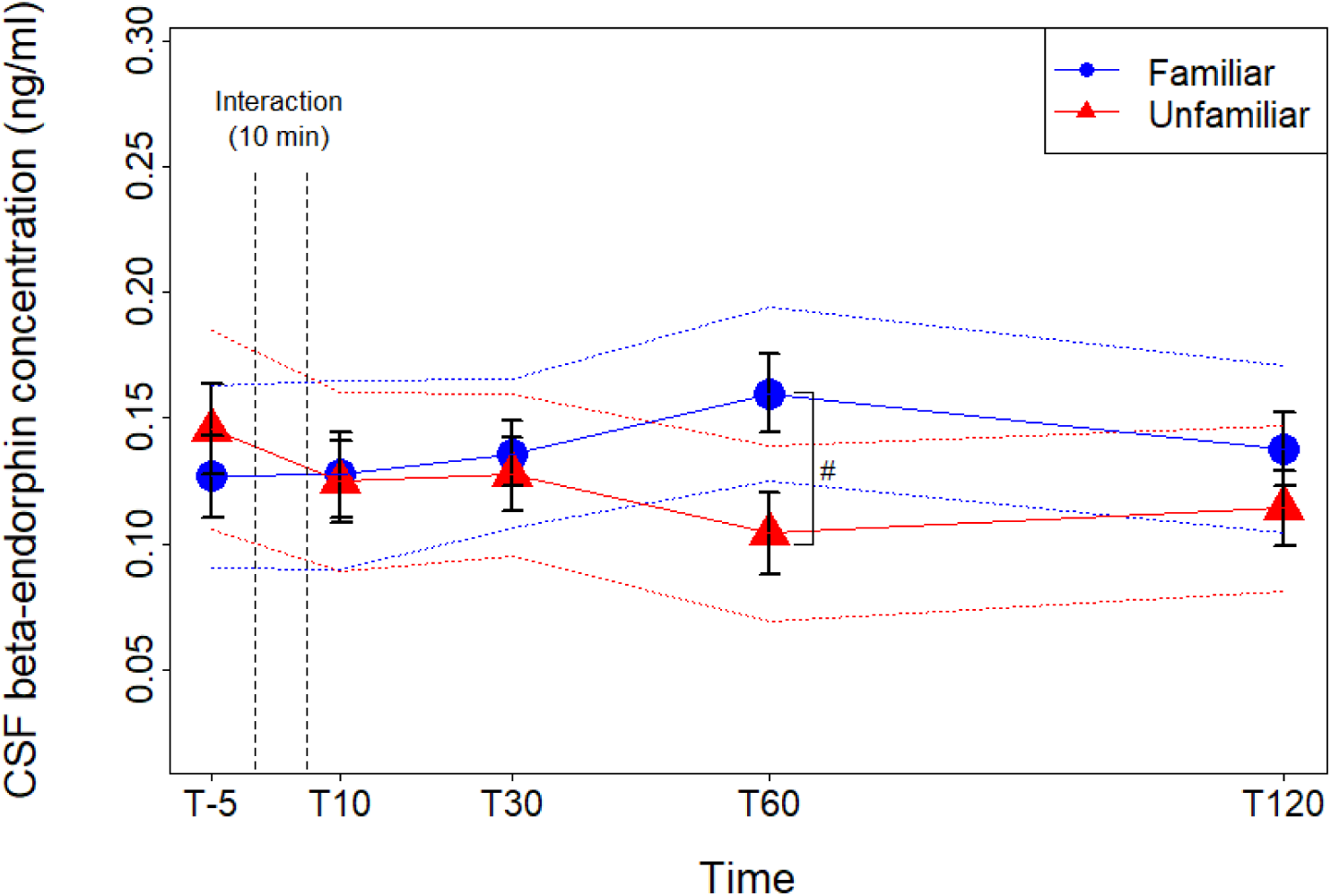
Effects of the human familiarity across times on the concentration of β-endorphin. The graph shows the means (dots/triangles) and standard errors (error bars), as well as the 95% confidence intervals (dotted lines) estimated from the statistical model. The statistical significance of the interaction between the human familiarity and time was P = 0.001, and the difference between the two human familiarities at T60 was a tendency (P = 0.055) represented by the symbol “#”.

### 3.5. Behaviour

None of the factors of interest (i.e. human familiarity, contact type, or their interaction) had a significant effect on the total duration of contact initiated by the pig or on the frequency of the contacts initiated by the pig (Table S1, Figure S6). However, there was a tendency for the frequency of contacts initiated by the pig to be higher with the “unfamiliar human” than with the “familiar human” (3.22 ± 0.45 vs 2.6 ± 0.33; t_15.18_ = 1.8; P = 0.09; Figure 6), independently of the contact type.

**Figure 6.**
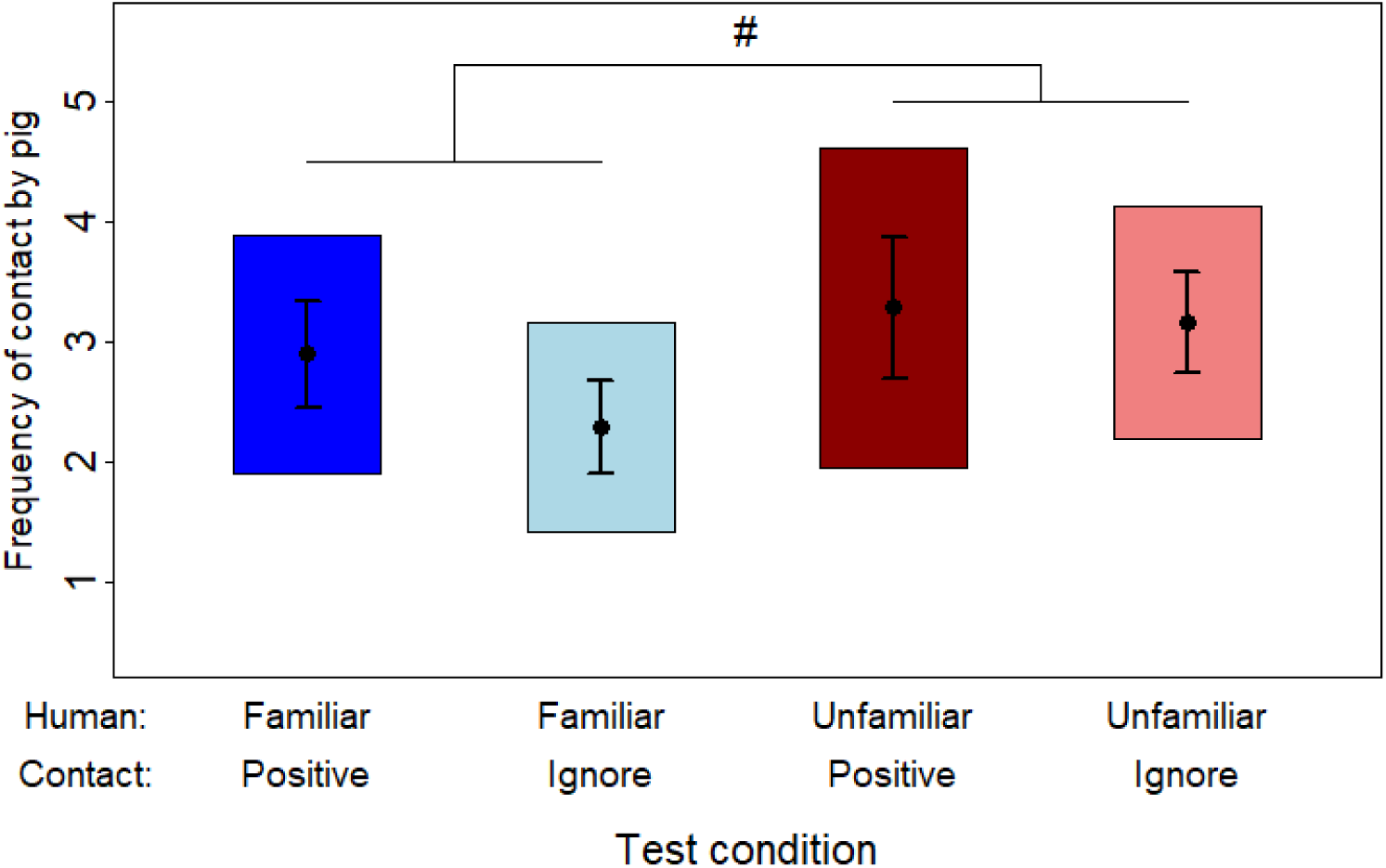
Overall effects of the test conditions (interaction between the human familiarity and the type of contacts) on the frequency of contacts initiated by the pig during the 10-min interaction period. The graph shows the means (dots) and standard errors (error bars), as well as the 95% confidence intervals (box) estimated from the statistical model. The tendency for difference between the human familiarity levels is represented by the symbol “#” (P = 0.091).

The effect of the behaviour on the concentration of neurotransmitters at the first timepoint post-test was investigated (i.e. T10 for 5-HIAA, DOPAC, HVA and β-END and T60 for OT). Longer total duration of contact initiated by the pig during the test session was linked to higher HVA concentration at T10 (estimate = 0.02 ± 0.005; t_30.73_ = 3.55; P = 0.001). The other neurotransmitters (OT, DOPAC, 5-HIAA and β-END) were not significantly linked to the behaviour of the pigs during the test.

### 3.6. Proteome

The analysis of the pig proteome (1619 genome wide quantified proteins) did not reveal any significant effect of the factors of interest (human familiarity, contact type, or their interaction) but showed large animal-specific effects. The heatmap (Figure 7) showed that proteins clustered column-wise according to animals. The violin plots of variances (Figure 8) revealed that animal specific effects explained the majority of the amount of variance across all 1619 proteins. In a mixed model, the random intercept component measures the variance associated with individual random pig effect deviations to the estimated fixed effect intercept, and it was the second most important source of variance in our proteomics data. Random slopes model explained the interaction between fixed effects and levels of random effect, i.e. pig specific deviations for each contrast of our 8 factor-level interaction between human familiarity, type of contact and time and interestingly the biggest source of variance in our proteomics data. There was very little technical noise (housing batch and date of protein quantification) and almost no variance associated with our treatment of interest by random intercept of pig. Furthermore, the seven dummy-coded centred random slopes exceeded the variance explained by our experimental variables of interest and their interaction and possible sources of technical noise, pigs’ batch and date of quantification, respectively.

**Figure 7.**
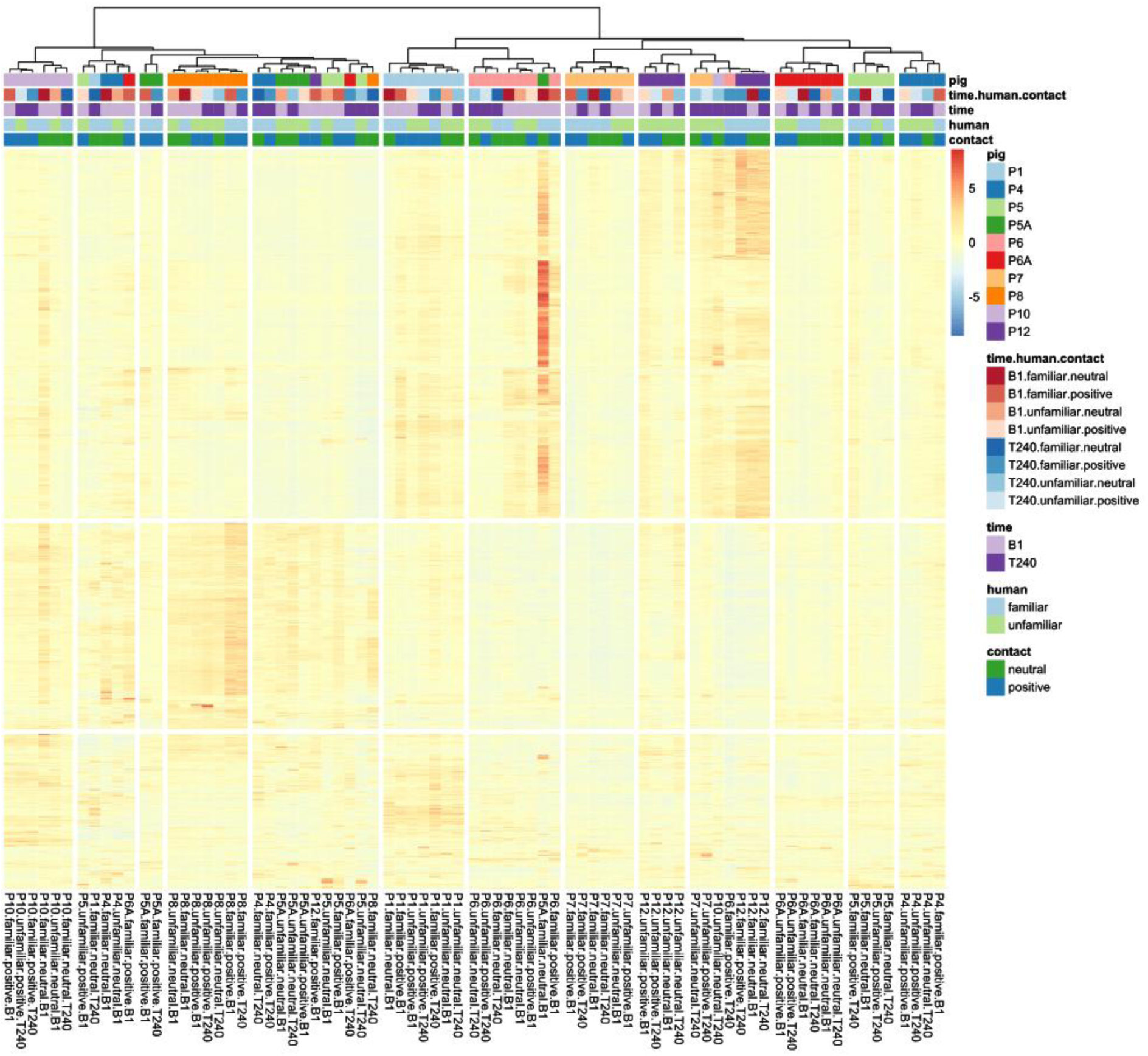
Unsupervised row-wise and column-wise clustering of 1619 quantified, row-wise z-transformed proteins, with correlation coefficients as distance measures, using Ward’s hierarchical agglomerative clustering method (Murtagh and Legendre, 2014). The heatmap is annotated with information on sample ID, pig, and the experimental variables of interest: time (pre-test (B1) measure and 240 minutes post-test (T240)), human familiarity (familiar and unfamiliar), contact type (ignore (“neutral”) or positive), and the combination of the above.

**Figure 8.**
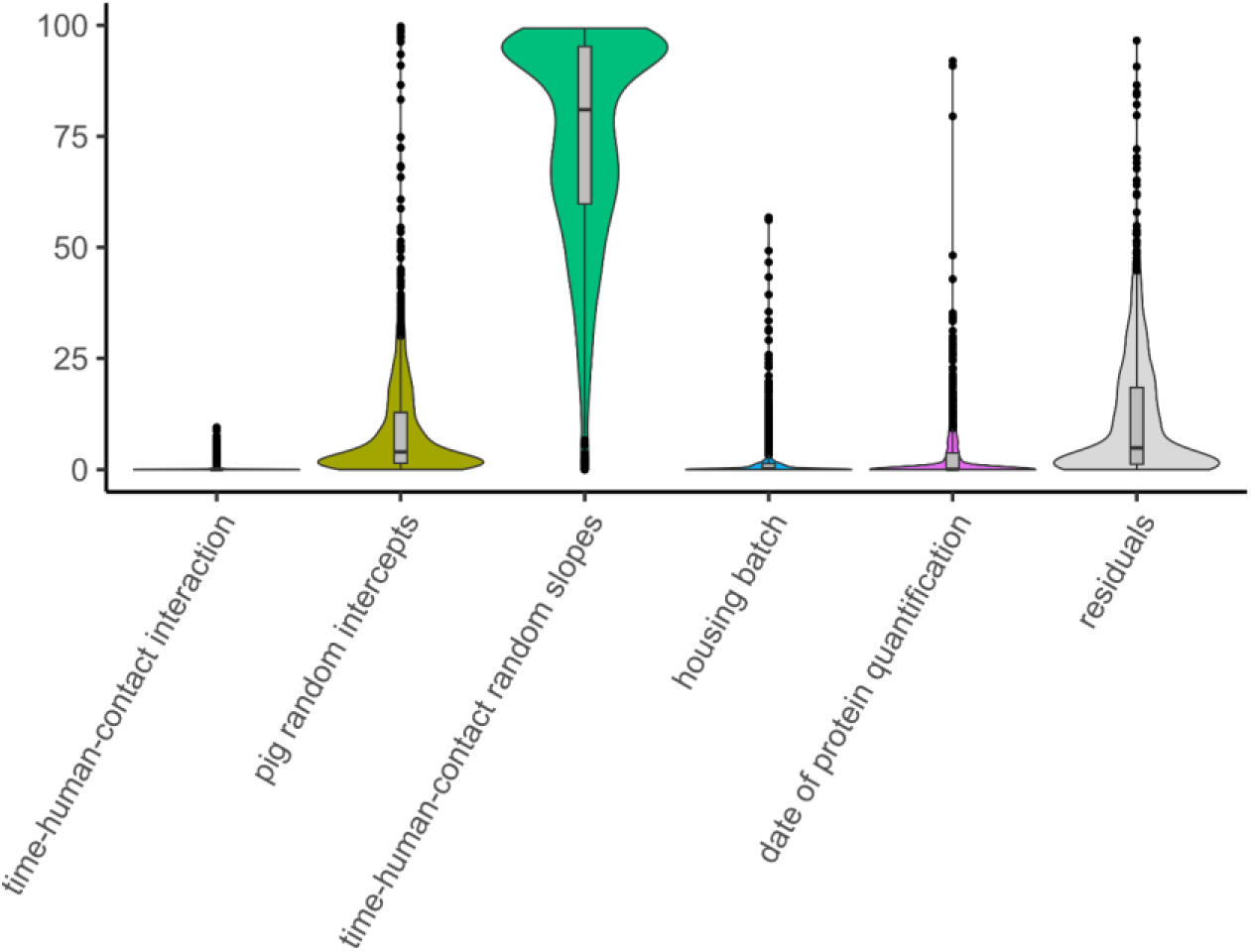
Violin plots of variances explained by simultaneously estimated random effects for each of 1619 log10 transformed quantified proteins, after adding a constant of 1 to each observation.

## 4. Discussion

This study aimed to assess the neurotransmitter dynamics induced by positive human-pig interactions. We hypothesised that greater concentrations of all neurotransmitters would be observed after a session involving a familiar human delivering positive contacts, as a reflection of the positive valence of such interaction for the pigs. Conversely, we hypothesised that the unfamiliar human ignoring the pigs will constitute a less pleasurable experience for the pigs and would lead to the lowest concentrations of neurotransmitters. However, our results contradicted these predictions.

Overall, the familiarity of the human interacting with the pigs (familiar or unfamiliar) did not significantly affect the neurotransmitter concentrations, apart from a tendency for β-END concentration to be higher 1 h after interacting with the familiar human than with the unfamiliar human. This supports the brain opioid theory of social attachment where endogenous opioids, particularly β-END, have been heavily implicated in social bonding (Machin and Dunbar, 2011) with some having shown that the effects can be mediated by the emotional closeness (i.e. stronger social bond) to the other (Inagaki et al., 2020).

Our results are contrary to our prediction that OT underlies social selectivity and would be higher after interacting with the familiar human compared to the unfamiliar human. Instead, we found the opposite, with the highest OT concentration when the unfamiliar human ignored the pig, which is in line with some of the literature that suggests that OT is released during encounters with unfamiliar individuals or when faced with a novel situation (Tops et al., 2013a). This, along with the tendency for pigs to initiate higher frequency of contacts toward the unfamiliar human, supports the theory that OT release facilitates approach behaviour (Heinrichs and Domes, 2008).

For all neurotransmitters studied except HVA, the interaction between human familiarity and contact type was statistically significant (5-HIAA, β-END) or a tendency (DOPAC, OT) but the pairwise comparisons were not significant (except for OT). This is likely due to the small sample size of the study, as the success rate of functioning spinal catheters was relatively low (44%, 16/36 pigs; 6 pigs were not included in the analysis due to incomplete dataset, i.e. missing samples in some conditions or at some timepoints). Furthermore, our study aimed to detect perhaps more subtle changes than comparable studies, as we investigated the effect of providing positive contacts compared to ignoring the pigs, which might lead to smaller differences than comparing positive to negative contacts (Boissy et al., 2007).

The absence of marked differences in the neurotransmitter concentrations and pig behaviour across the test situations suggests that the interactions might have been overall considered positive by the pigs (in 60% of sessions the pigs spent 50-99% of the time in contact with the human). Indeed, the higher concentration of HVA at T30 compared to pre-test, together with the link between HVA concentration and duration of contact by the pig, irrespective of treatment condition, suggest that the pigs were motivated to interact or find it rewarding regardless of the familiarity of the human or type of interaction. The duration of contact could be related to investigation or exploratory behaviour driven by dopamine (Kesner et al., 2022), albeit this association failed to be demonstrated in monkeys (Parker et al., 2007), also as pigs initiated more frequent contact toward the unfamiliar human than the familiar human. Because the pigs were only exposed to positive contacts during habituation (interacting human, handler) and gentle interactions from all caretakers, they probably generalised their experience to all humans (Brajon et al., 2015), including the unfamiliar humans. In addition, the unfamiliar humans were instructed to interact in a similar way to the familiar human to minimise differences in interaction style, which may have minimized differences. Furthermore, the pigs were still able to interact with the human during the ignoring condition as they pleased (e.g. nudging, chewing on overall, lying in contact), and this might have been as rewarding and pleasurable for them as when the human was actively delivering positive contacts. Although not analysed statistically, pigs showed belly rubbing in 14 out of 23 sessions involving positive contacts, of which 10 were with the familiar human and 4 were with the unfamiliar human. Belly-rubbing is a behaviour that sometimes occur during positive interactions, during which the pigs lie laterally and allow the human to rub their belly while grunting softly and closing their eyes (Rault et al., 2019). This type of behaviour is thought to be indicative of positive welfare state and might reflect trust (Rault et al., 2019), suggesting that the pigs might have perceived all interactions as positive but still made a distinction between the unfamiliar and familiar humans.

There was no clustering of the pigs’ proteome according to the test conditions, i.e. the identity of human, the type of contact and their combination, but there was a strong individual animal clustering. Indeed, the large inter-individual variation in the proteome observed made it difficult to identify treatment effects with such a small sample size. Furthermore, the timing of sampling (240 min post-interaction) might have been too long after the interaction to allow for the identification of changes in the proteome in relation to the interaction. However, the comparison of the clustering displayed in the heatmaps and violin plots between different models run suggests that the variances can indeed be attributed to individual variation rather than technical noise, i.e. the residual variances are much smaller when fitting animal effects instead of injection times during proteomics analysis. This part of the study was explorative, given the novelty of proteomics studies in general and the absence of comparable studies. However, our results suggest that investigations of proteome changes in relationship with human-animal relationship require large sample sizes to be able to detect the subtle effects of such treatments. Furthermore, studies should consider collecting samples closer to the time of the interaction.

Finally, some aspects of the methodology used should be discussed. First of all, the lack of reference for CSF sampling timepoints to observe changes in β-END concentration and proteomics made our investigation quite exploratory, and it is probable that the timing chosen for sampling timepoints were not optimal to detect changes. Similarly, it is unknown whether the sampling timepoints chosen in this study were able to capture transient changes in DA and 5-HT given these usually occur on a sub-second to second timescale (Yagishita, 2020), or long enough to capture 5-HT’s effects during the consummatory phase (Liu et al., 2020). Secondly, although it was originally planned to obtain CSF samples from 36 pigs, there was a large dropout rate (i.e. catheter placement failure: 4 pigs, non-functional catheters: 16 pigs, partly functional catheters: 6 pigs) and practical reasons prevented the recruitment of an additional batch of pigs. A potential cause for non-functional catheter was its misplacement, *i.e.* outside the subarachnoid space (as determined by post-mortem fluoroscopy in some pigs; data not shown), but for most of the pigs the reason of failure to collect CSF could not be identified. Therefore, despite the obvious advantages of using CSF sampling methods to capture changes in neurotransmitters dynamics at the central level, the methodology of spinal catheter placement needs further refinement to increase success rate.

In conclusion, subtle differences in pigs’ neuroendocrine responses to human-animal interactions were detected, although these were likely overshadowed by large individual variation and the small sample size. Opioids rather than oxytocin differed according to familiarity, supporting the brain opioid theory of social attachment. The increased oxytocin concentration along with frequency of contacts initiated by the pig toward an unfamiliar human ignoring the pig supports the literature implicating oxytocin’s role in facilitating approach and bonding with unfamiliar individuals or more generally being released in novel situations. The lack of clear differences between our four treatment conditions (human familiarity × contact type) suggests that the pigs perceived all interactions to be overall positive, perhaps due to generalisation of their positive experiences throughout the habituation period, and because they were still free to interact with humans whilst the human was ignoring them. This is the first study to measure oxytocin, dopamine, opioids and serotonin concurrently, which shows the promise of this approach to advance the understanding of the neurophysiological basis of positive interactions.

## Acknowledgements

We thank the staff at the Medau farm and at the Swine Clinic of the University of Veterinary Medicine, Vienna for taking care of the pigs and providing help in the conduction of this experiment. We would like to address a special thank you to the students who acted as the unfamiliar human in the tests (Edie Challinor, Asta Proksch) and who helped with their care (Maiwen Braconnier). We would also like to thank Dagmar Mähling from the FBN Dummerstorf for her technical support in analysing monoamines in CSF samples; and André Müller, Thomas Hannich and Anna Tinnacher from the Research Center for Molecular Medicine of the Austrian Academy of Sciences (CeMM) for processing our CSF samples for proteomics analysis.

## Authors contributions

Conceptualization: OS, JLR, ST

Data curation: OS

Formal analysis: OS, WO, HV, KH, MD,

Funding acquisition: JLR

Investigation: OS, ST, JG, AR

Methodology: OS, ST, JLR

Project administration: OS

Resources: WO, AR

Supervision: JLR

Visualization: OS, MD

Writing – original draft: OS, ST

Writing – review and editing: OS, ST, JLR, WO, MD, KH, JG, HV, AR

