## Supplementary materials for "The inner dynamics of positive human-animal interactions: investigating the roles of oxytocin, opioids, dopamine, serotonin and the proteome"

### Supplementary material

**Table S1.** Ethogram for behavioural observation of the test sessions. Behaviours within categories are mutually exclusive unless stated otherwise. Interruptions shorter than 3s were considered the same bout of behaviour.

| Category | Behaviour | Modifier | Definition |
|---|---|---|---|
| *Posture* | | | |
| | Standing | | Pig's torso does not touch the floor, standing on four legs. |
| | Sitting | | Pig's hind legs folded underneath the body and supporting weight on the two front legs. "Sitting like a dog" |
| | Lying | Lying on sternum, Lying laterally | Pig's torso touches the floor, lying on the belly or on the side. |
| *Activity* | | | |
| | Contact by pig | | Pig makes physical contact with human. Includes active behaviours such as sniffing, nosing, oral manipulation, rubbing, pawing, and passive behaviours such as standing, sitting or lying in physical contact with human's body. |
| | Contact by human[a,b] | Belly rubbing, All other contact | Pig receives physical contact by the human. Includes active contact (stroking, scratching, rubbing) and passive hand resting on the body of the pig. |
| | Physical interaction[b] | | Pig and human are in physical contact with each other. |
| | Locomotion | | Walking or running; lifting and setting down two or more feet to form one or more steps. |
| | Exploring the pen | | Pig's snout touches pen floor or wall for more than 1s. |
| | Immobile | Look at exit door, Look elsewhere | Pig is either standing, sitting or lying for more than 2s and less than 30s, shows no specific movement and is not engaged in any other activity. |
| | Inactive | | Pig is either standing or lying, shows no specific movement for more than 30 seconds. The pig remains inactive if it engages in exploration maximum twice or for less than 5s, takes less than five steps away, stretches or changes posture. |
| | Look for escape | | Pig nudges/pushes against pen door using snout or front legs; climbs door or wall; pacing around pen and looking up. |

| | | | |
|---|---|---|---|
| | Escape attempt[c] | | Pig jumps against the door or wall. |
| | Elimination[c] | | Pig eliminates faeces or excretes urine. |
| | Other[b] | | Any other behaviour not listed above. Make a note. |
| | Not visible | With human, Without human | Pig's behaviour cannot be determined. |
| *Tail posture and movement* | | | |
| | Hanging tail | | Tail hanging down against the body. |
| | Curled tail | | Tail coiled up in a curl on top of the body. |
| | Tucked tail | | Pig's tail is kept vertically down and close to the body whereby the tip of the tail is held between the legs. |
| | Wagging | | Tail swinging in any direction, but mostly from side to side. |
| | Not visible | | Pig's tail posture or movement cannot be determined. |

[a] Familiar positive and Unfamiliar positive conditions only.
[b] Behaviour does not exclude any other behaviours.
[c] Scored as point event due to short duration.

**Table S2.** Summary of the output (mean ± S.E.) estimated by the statistical models

| | | Positive contacts | | Neutral contacts (ignore) | | | | | | | | |
|---|---|---|---|---|---|---|---|---|---|---|---|---|
| | | **Familiar human** | **Unfamiliar human** | **Familiar human** | **Unfamiliar human** | Effect contact | Effect human | Effect contact × human | Effect time | Effect time × contact | Effect time × human | Effect contact × human × time |
| Oxytocin (pg/ml) | T-60 | 15.1 ± 2.02 | 15.0 ± 2.02 | 20.8 ± 2.09 | 18.8 ± 1.98 | 0.01 | 0.49 | 0.08 | 0.00 | 0.37 | 0.44 | 0.26 |
| | T60 | 19.3 ± 2.22 | 18.1 ± 2.22 | 20.4 ± 2.17 | 21.8 ± 2.07 | | | | | | | |
| | T120 | 18.4 ± 2.52 | 17.7 ± 2.61 | 19.0 ± 2.46 | 22.2 ± 2.46 | | | | | | | |
| | T240 | 16.9 ± 1.98 | 10.4 ± 1.98 | 13.5 ± 1.95 | 14.7 ± 2.07 | | | | | | | |
| DOPAC (ng/ml) | T-10 | 7.2 ± 0.74 | 6.9 ± 1.20 | 7.8 ± 1.50 | 6.1 ± 1.68 | 0.96 | 0.53 | 0.07 | 0.80 | 0.43 | 0.56 | 0.98 |
| | T10 | 6.7 ± 0.66 | 7.1 ± 1.11 | 8.1 ± 1.50 | 7.1 ± 1.69 | | | | | | | |
| | T30 | 7.2 ± 0.75 | 7.5 ± 1.16 | 7.4 ± 1.38 | 6.6 ± 1.54 | | | | | | | |
| HVA (ng/ml) | T-10 | 22.8 ± 2.64 | 21.8 ± 4.29 | 23.2 ± 3.28 | 25.7 ± 4.81 | 0.83 | 0.38 | 0.92 | 0.02 | 0.44 | 0.56 | 0.44 |
| | T10 | 25.0 ± 2.23 | 27.5 ± 3.79 | 24.9 ± 2.98 | 26.2 ± 4.48 | | | | | | | |
| | T30 | 25.0 ± 2.84 | 29.2 ± 4.40 | 25.9 ± 3.31 | 28.2 ± 4.79 | | | | | | | |
| 5-HIAA (ng/ml) | T-10 | 15.7 ± 1.34 | 16.8 ± 1.60 | 16.6 ± 1.42 | 16.5 ± 1.50 | 0.80 | 0.47 | 0.03 | 0.13 | 0.99 | 0.79 | 0.73 |
| | T10 | 15.5 ± 1.28 | 17.8 ± 1.53 | 17.2 ± 1.46 | 16.6 ± 1.60 | | | | | | | |
| | T30 | 16.1 ± 1.51 | 18.8 ± 1.73 | 17.9 ± 1.44 | 17.8 ± 1.49 | | | | | | | |
| β-endorphin (ng/ml) | T-10 | 0.13 ± 0.02 | 0.13 ± 0.02 | 0.12 ± 0.02 | 0.16 ± 0.02 | 0.45 | 0.25 | 0.04 | 0.84 | 0.83 | 0.00 | 0.60 |
| | T10 | 0.12 ± 0.02 | 0.12 ± 0.02 | 0.14 ± 0.02 | 0.13 ± 0.02 | | | | | | | |
| | T30 | 0.14 ± 0.02 | 0.12 ± 0.02 | 0.13 ± 0.02 | 0.13 ± 0.02 | | | | | | | |
| | T60 | 0.16 ± 0.02 | 0.09 ± 0.02 | 0.16 ± 0.02 | 0.12 ± 0.02 | | | | | | | |
| | T120 | 0.15 ± 0.02 | 0.11 ± 0.02 | 0.13 ± 0.02 | 0.12 ± 0.02 | | | | | | | |
| Contact by pig (min) | | 380 ± 55.0 | 410 ± 41.8 | 345 ± 52.0 | 322 ± 39.7 | 0.18 | 0.74 | 0.44 | | | | |
| Contact by pig (n/min) | | 2.9 ± 0.44 | 3.3 ± 0.58 | 3.85 ± 0.39 | 3.16 ± 0.42 | 0.37 | 0.09 | 0.417 | | | | |

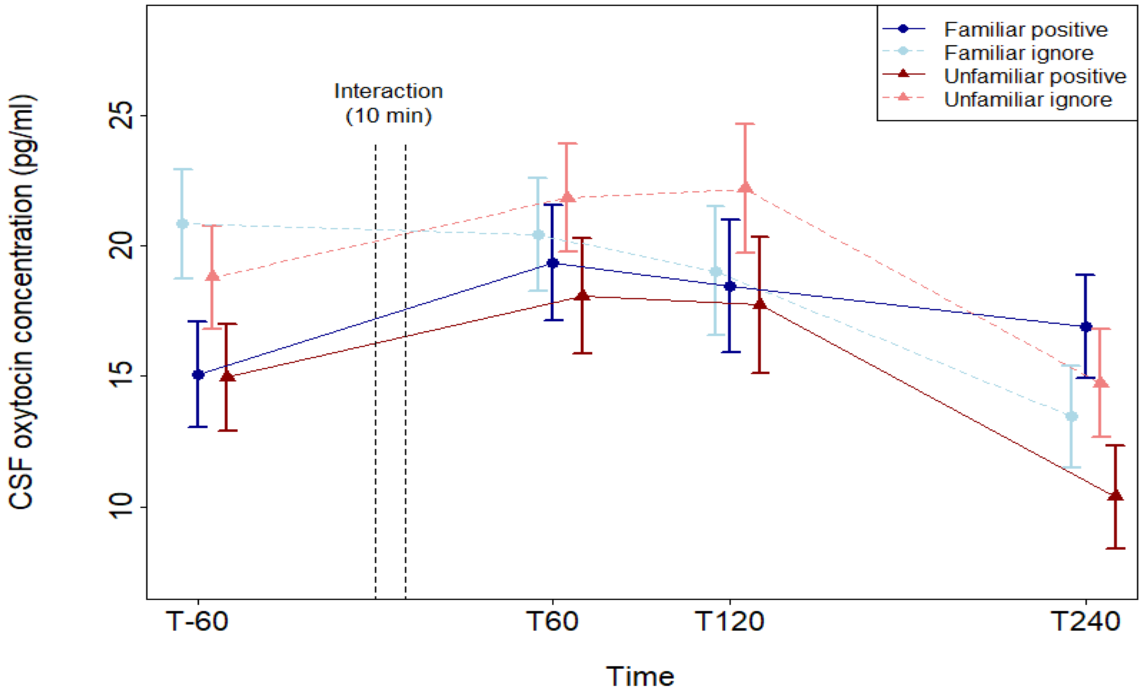


**Figure S1.** Mean and standard errors of the concentration of oxytocin in the cerebrospinal fluid (CSF) of pigs across time and according to the test conditions (interaction between human familiarity and contact type). Familiar human = dots, Unfamiliar human = triangles, positive contacts = full lines, neutral contacts (neutral) = dotted lines.

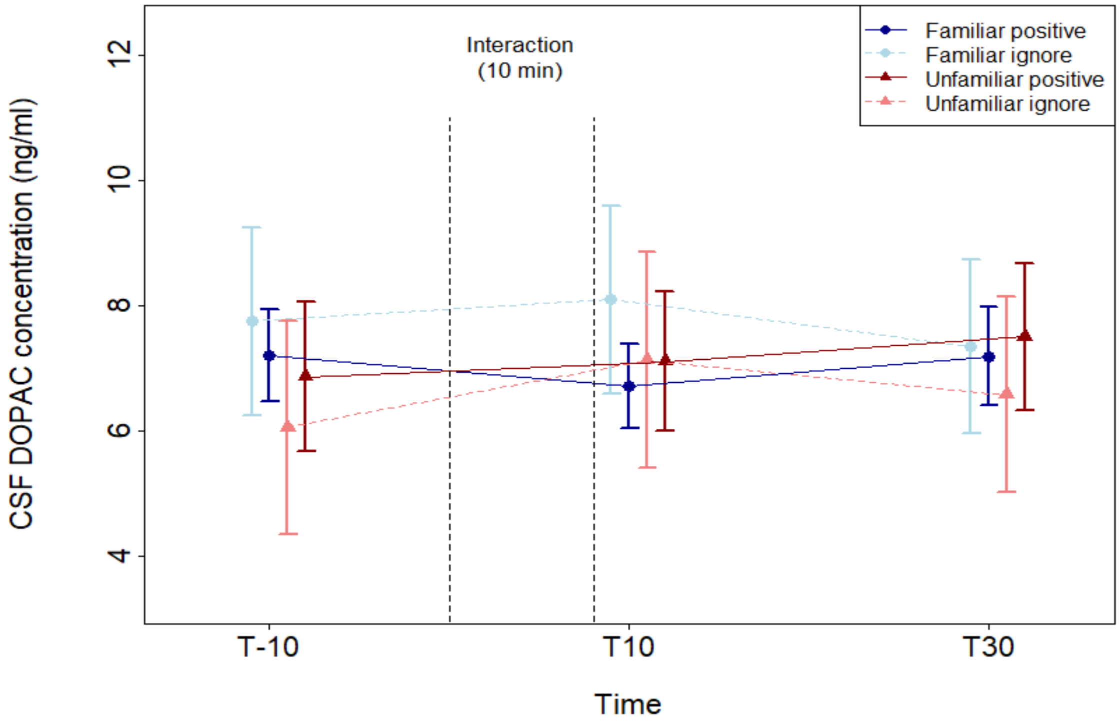


**Figure S2.** Mean and standard errors of the concentration of DOPAC in the cerebrospinal fluid (CSF) of pigs across time and according to the test conditions (interaction between human familiarity and contact type). Familiar human = dots, Unfamiliar human = triangles, positive contacts = full lines, neutral contacts (neutral) = dotted lines.

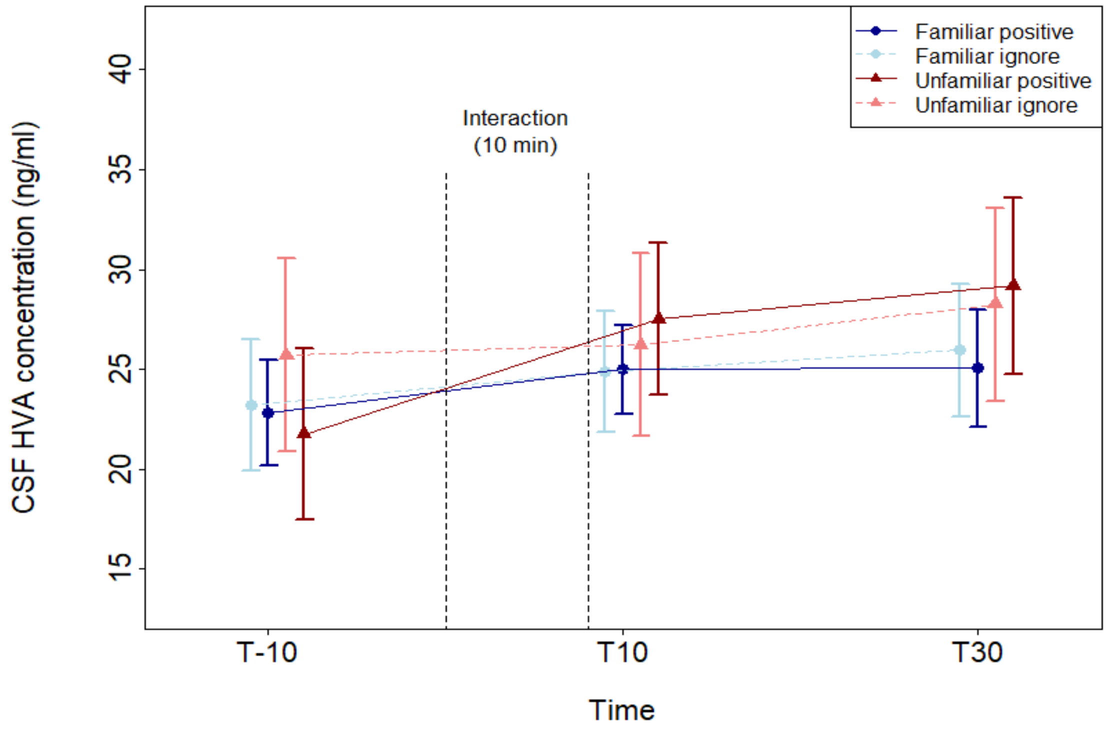


**Figure S3.** Mean and standard errors of the concentration of HVA in the cerebrospinal fluid (CSF) of pigs across time and according to the test conditions (interaction between human familiarity and contact type). Familiar human = dots, Unfamiliar human = triangles, positive contacts = full lines, neutral contacts (neutral) = dotted lines.

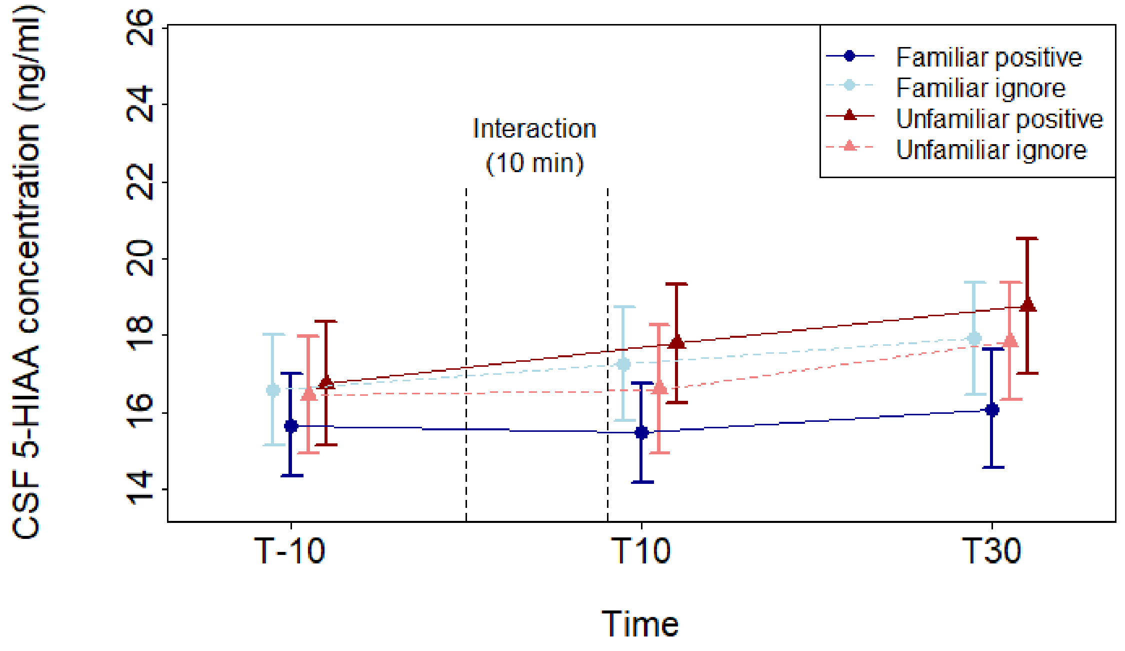


**Figure S4.** Mean and standard errors of the concentration of 5-HIAA in the cerebrospinal fluid (CSF) of pigs across time and according to the test conditions (interaction between human familiarity and contact type). Familiar human = dots, Unfamiliar human = triangles, positive contacts = full lines, neutral contacts (neutral) = dotted lines.

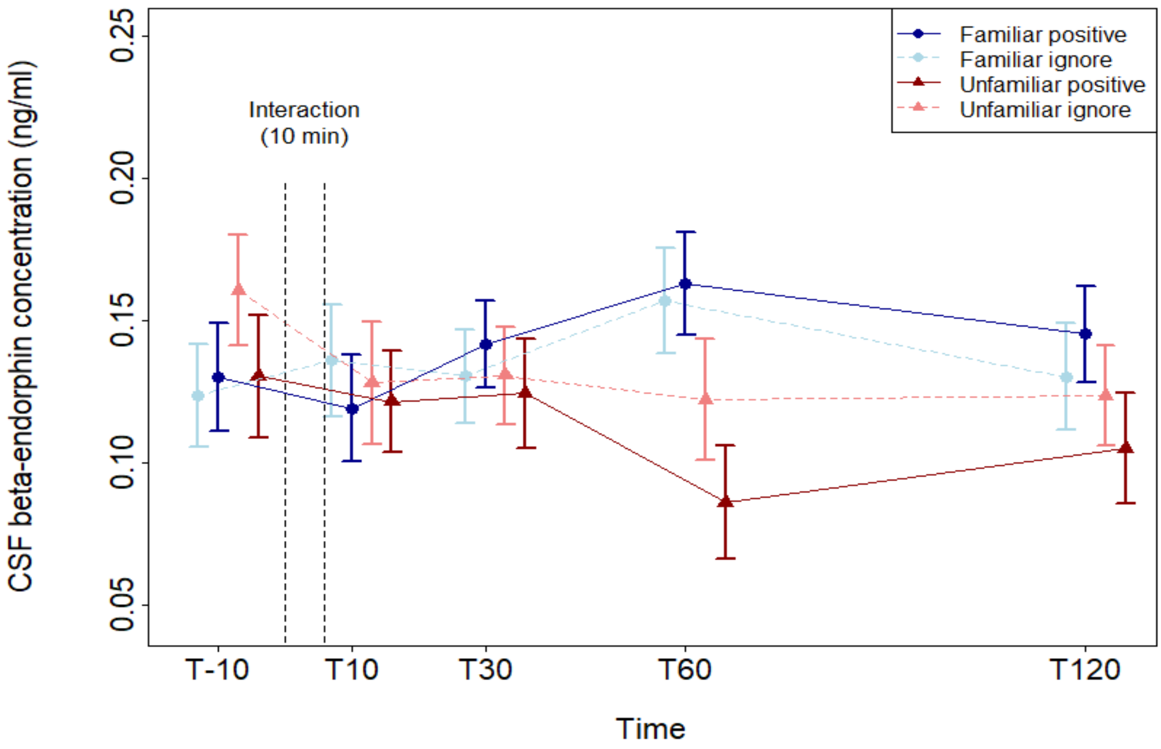


**Figure S5.** Mean and standard errors of the concentration of β-endorphin in the cerebrospinal fluid (CSF) of pigs across time and according to the test conditions (interaction between human familiarity and contact type). Familiar human = dots, Unfamiliar human = triangles, positive contacts = full lines, neutral contacts (neutral) = dotted lines.

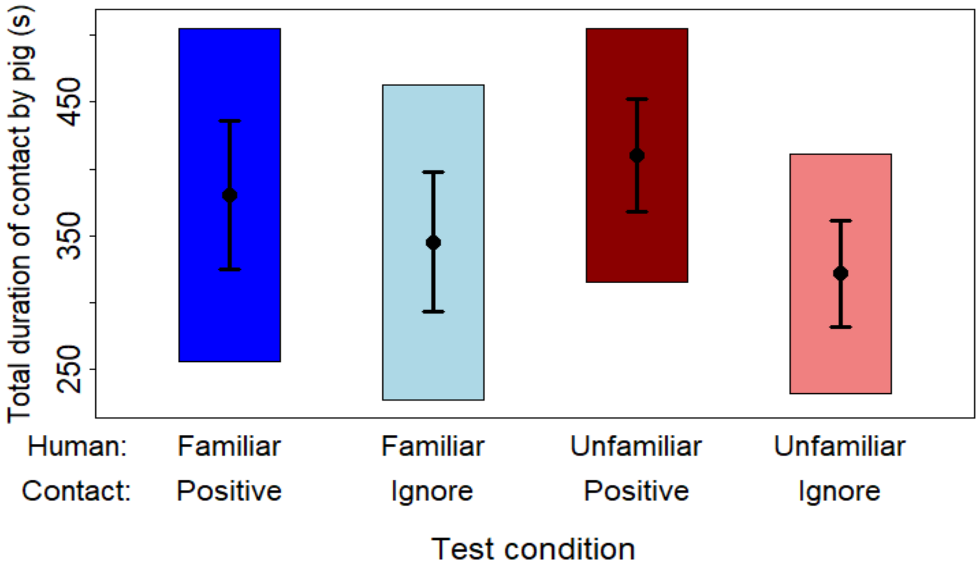


**Figure S6.** Overall effects of the test conditions (interaction between the human identity and the contacts type) on the duration of contacts initiated by the pig during the 10-min test sessions. The graph shows the means (dots) and standard errors (error bars), as well as the confidence limits (box) estimated from the statistical model.
